# Rainfall legacy effects on the rhizosphere bacterial diversity of *Brachypodium* ecotypes across an aridity gradient

**DOI:** 10.64898/2026.01.27.702010

**Authors:** Sheetal, Peter Reuven, Sofia Maite Arellano, Maor Matzrafi, Shilo Rosenwaser, Daniella Gat, Elisa Korenblum

**Affiliations:** Department of Plant Sciences, The Robert H. Smith Faculty of Agriculture, Food and Environment, Hebrew University of Jerusalem, Rehovot, Israel; Plant Sciences Institute, Agricultural Research Organization – Volcani Institute, Rishon LeZion, Israel; Department of Plant Pathology and Weed Research, Newe Ya’ar Research Center, Agricultural Research Organization (ARO) – Volcani Institute, Ramat Yishay, Israel

**Keywords:** Rhizosphere microbiome, *Brachypodium* ecotypes, aridity gradient, precipitation legacy, common garden experiment

## Abstract

Plants exhibit clinal trait variation along aridity gradients driven by strong selective pressures, yet whether plant–microbiome associations follow similar patterns remains unclear. *Brachypodium* spp., a model for temperate cereals spanning environments from hyper-arid to humid, provides an ideal system to test this hypothesis. Here, we compare rhizosphere bacterial communities of *Brachypodium* growing across arid, semi-arid, and dry sub-humid zones in Israel with those of ecotypes collected along the same precipitation gradient and grown under common-garden conditions. Rhizosphere bacterial diversity was highest in plants from mid-precipitation sites within the Mediterranean semi-arid transition zone, where annual precipitation decreases from 600 to 400 mm. Together with higher stochasticity and fewer significantly associated taxa in rhizosphere microbiomes from mid-precipitation plants suggest weaker plant-driven microbiome selection in this transition zone, a pattern that persisted under common-garden conditions. The results may represent a promising avenue to develop microbiome-based strategies for drought resilience by advancing our understanding of host filtering across aridity gradients.

## Introduction

The rhizosphere microbiome is one of the most complex and biologically active ecosystems on Earth, harbouring an estimated 10^8^ microbial cells per gram of soil (Whitman et al., 1998). This specialized region plays a critical role in plant health, growth, defence mechanisms and nutrient regulation (Mueller & Sachs, 2015). The rhizosphere contains various microorganisms (e.g., bacteria, fungi, archaea, oomycetes, and protists) that collectively influence plant development and resistance to both biotic and abiotic stressors. The rhizosphere microbial composition is dominated by prokaryotic phyla including Pseudomonadota, Actinomycetota, Bacteroidota, Bacillota and Acidobacteriota (Li et al., 2024). The composition of microbial communities is affected by various environmental factors such as pH, precipitation, soil texture and chemistry, temperature (Fierer et al., 2003; Fierer & Jackson, 2006; Ramirez et al., 2012; Tripathi et al., 2017); as well as biological factors, including trophic interactions and plant-derived compounds (Compant et al., 2019). Since plants often depend on their root-associated microbiota to cope with environmental stress, shifts in the composition and function of the root microbiome due to external factors can undermine its beneficial effects on plants (Vandenkoornhuyse et al., 2015), particularly under water-limited conditions. Precipitation variation was associated with changes in soil microbial communities in various habitats of desert ephemeral plants (Tripathi et al., 2017). In drylands worldwide, aridity drives compositional shifts, favoring drought-tolerant taxa such as Chloroflexi and α- Proteobacteria while reducing Acidobacteriota and Verrucomicrobiota and reduces soil microbial diversity and abundance.

Microbiome composition shifts threaten essential ecosystem services, suggesting that co-adaptation between plants and microbes will be increasingly critical for maintaining soil fertility and resilience as drylands expand under climate change (Maestre et al., 2015). Drought selective pressure favours microbial taxa in the rhizosphere that enhance drought resilience through osmolyte production, phytohormone modulation, and improved nutrient cycling. Moreover, shifts in temperature and moisture regimes modify root traits and exudation patterns, influencing microbial community assembly and functional diversity. Thus, drylands are among the most extreme environments, characterized by low and variable rainfall, where water availability influences both plant growth and microbial assembly (Pugnaire et al., 2019).

As aridity increases, plants often exhibit smaller, thicker leaves to reduce transpiration, deeper or more extensive root systems for improved water uptake, and altered stomatal density to optimize gas exchange under drought stress. Trait adjustment across gradients (clinal variation) indicates strong selective pressures and has been largely studied across physiological responses, growth, genomic features, and life-history traits (Adams et al., 2016). However, the extent to which clinal variation in plants is reflected in plant-microbiome associations remains largely unknown. Recent works show that conditions of the habitat of origin explain the microbiome variation of different plant ecotypes (Alegria Terrazas et al., 2020; Ricks et al., 2026). Consequently, co-adaptation along aridity gradients emerges as a critical mechanism for maintaining plant resilience and ecosystem functioning.

*Brachypodium* spp., a model species for cereal crops such as wheat, naturally occurs in hyper-arid to humid habitats and may possess adaptive strategies mediated through its rhizosphere microbiome. According to recent results by Ricks et al., (2026), *Brachypodium* plants grown in a common garden display a rhizosphere microbiome that echoes the environmental conditions in their habitat of origin. Here, we investigated the rhizosphere microbiome of *Brachypodium* plants naturally growing in various locations in Israel, representing three different climate zones (arid, semi-arid, and dry sub-humid), and compared the bacterial diversities to the rhizosphere microbiomes of *Brachypodium* ecotypes previously collected from different locations in Israel (Penner et al., 2020) and grown from seeds in a common garden.

## Methods

### 1. Natural population (NP) sample collection

To investigate the rhizosphere microbiome of *Brachypodium* spp., plant root samples were collected from seven different locations across Israel in April 2022 (NP, Fig. 1), from three to four plants per location, all in their flowering stage. All soil particles attached to the roots were gently removed, and the collected roots were stored in sterile 15 ml tubes and kept on dry ice until transferred to –80°C freezer, awaiting DNA extraction. From each plant, a leaf was collected to a 1.5 ml tube to allow species identification of each plant. The seven locations of the sample collection are listed in table S1. Precipitation data for NP were obtained from the Israel Meteorological Service (IMS) database. Annual precipitation values for NP were calculated as the 5-year average preceding the time of collection (January 2017 – December 2021). These locations capture three different climate zones and span a precipitation gradient from ca. 300 to 700 mm per year.

**Figure 1.**
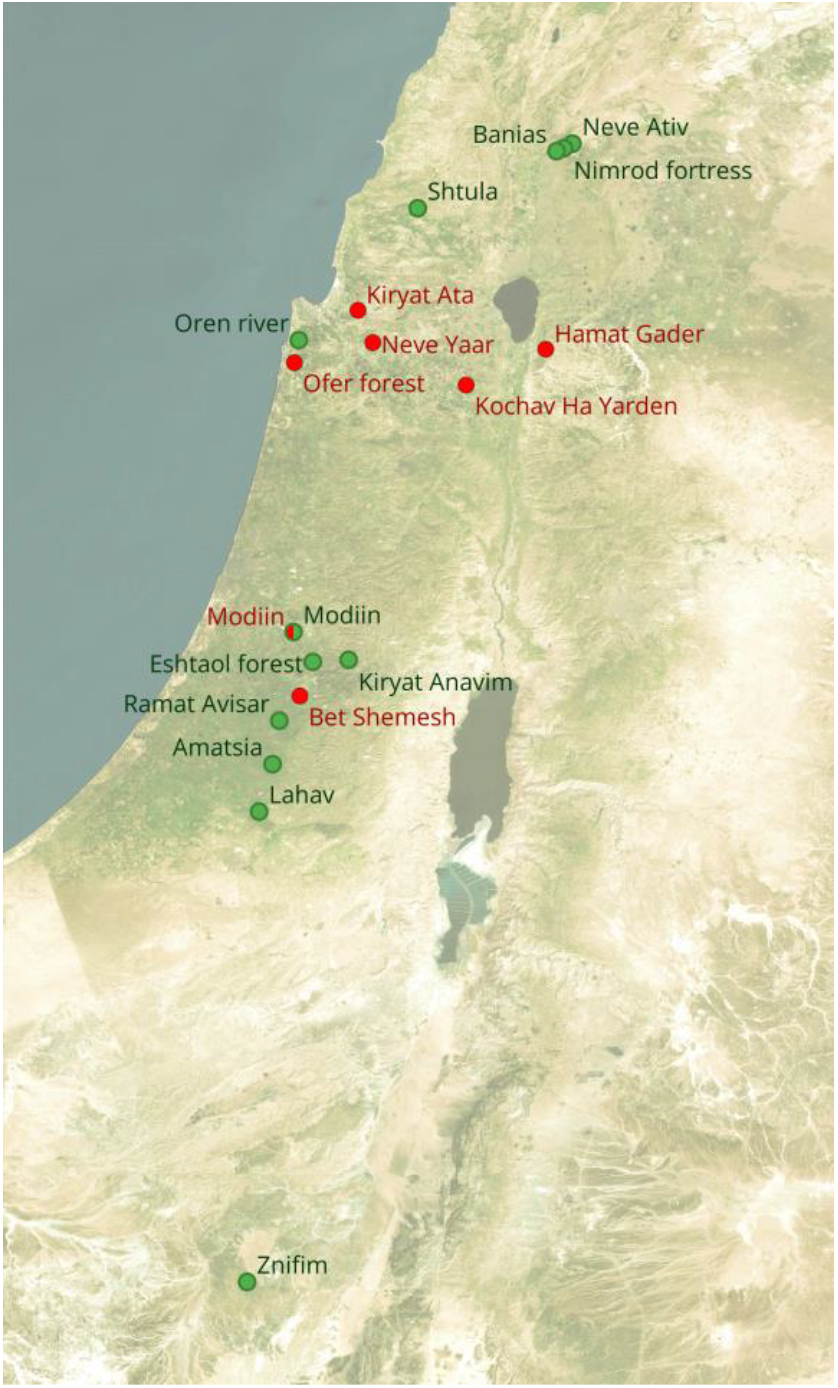
Geographical distribution of *Brachypodium* spp. Sampling sites across Israel. Map showing the locations of NP samples (indicated in red) and CGP samples (indicated in green) collected for this study was made using QGIS 3.17 based on the provided coordinates of the sampling points (QGIS Development Team, 2026).

### 2. Common garden (CGP) experiment

For the common garden experiment (CGP), we received *Brachypodium* seeds from Prof. Yuval Sapir (Tel-Aviv University, Israel, (Penner et al., 2020)). These seeds were collected from plants that were grown for at least two generations in a common Garden, under the same set of conditions, and originated from seeds of plans that were collected from 12 different locations in Israel (Table S1). For our common garden experiment, we planted these seeds in loam soil (sand - 37-65%; silt - 29-19% and clay - 34-16%) and incubated them in a controlled growth room under identical conditions with consistent watering and day light time of 16 hours, temperature 20°C. Weeds that grew in the pots were immediately removed manually. The plants were grown until flowering; their roots were collected similarly to the NP experiment. All samples and environmental data are presented in Table S1. Mean annual precipitation values for CGP were obtained from (Penner et al., 2020).

### 3. DNA extraction, amplification and sequencing

DNA was extracted from the rhizosphere of both NP and CGP samples using in-house phenol-chloroform method (Gat et al., 2025). PCR amplification of the 16S rRNA gene (a proxy for bacterial taxonomy) was conducted using the primer set 515F-806R (Caporaso et al., 2011). Each forward primer was barcoded with a unique 5-mer sequence to allow pooling of samples prior to library preparation. The reaction was conducted using 12.5 μl MyTaq HS mix (Meridian Bioscience, USA), under the following thermal cycle conditions: 5 min at 95°C followed by 27 cycles of 30 s at 95°C, 30 s at 50°C, 2 min at 72°C, and a final elongation step at 72°C for 10 min. All PCR products were run on an agarose gel (1.2%, w/v) to examine the amplicons for size and amount. PCR products were pooled and cleaned using Wizard SV gel and PCR clean-up system (Promega, USA). DNA quality was assessed using TapeStation (Agilent, USA). Following adapter ligation, the libraries were amplified for an additional 14 cycles and sequenced using a MiSeq System (Illumina, USA, V2, 500 cycles, paired-end) at the Crown Institute for Genomics, G-INCPM facility at the Weizmann Institute of Science, Rehovot, Israel.

For *Brachypodium* ploidy characterization, we extracted DNA from leaves of NP and CGP samples using Invisorb® plant DNA extraction kit, according to manufacturer instructions. We conducted PCR amplification using Primers F-ALB 16s and R-ALB 16s, followed by agarose gel electrophoresis, as described by (Giraldo et al., 2012). PCR amplification was carried out under the following cycling conditions: an initial denaturation step at 94°C for 2 minutes, followed by 40 cycles of 1 minute at 94°C, 1 minute at 56°C, and 1 minute at 72°C. A final extension step was performed at 72 °C for 10 minutes. PCR products were visualized by electrophoresis on a 2% agarose gel run at 100 mV for 60 minutes.

### 4. Sequence processing

The 16S rDNA sequences were demultiplexed using cutadapt v.4.5 and processed using DADA2 (v.1.26, (Callahan et al., 2016)) according to the suggested pipeline. Taxonomy assignment was conducted using the SILVA nr99, v.138_2 dataset. Mitochondria and plastid sequences, as well as non-bacteria and unclassified phyla, were manually removed. In addition, amplicon sequence variants (ASV) of the lowest 5-percentile read counts abundance and with a prevalence lower than 5% of the samples were removed to decrease sparsity. To correct for uneven sequencing depth, ASV counts were rarefied to the size of the smallest sample: 4856 and 8522 reads per sample for NP and CGP, respectively. Rarefaction was iterated 5,000 times and 10,000 times for NP and CGP ASVs, respectively (rarefaction curves presented in Figure S1), and the mean count per ASV per sample was calculated (Schloss, 2024). The rarefied tables were used to calculate all alpha-diversity indices and UniFrac weighted distances. The rarefied counts were transformed using centred log-ratio (CLR) transformation, preceded by zero-replacement via the Geometric Bayesian multiplicative method implemented in the zCompositions package (v1.4.1), to address the compositional nature of the data (Gloor et al., 2017), prior to Euclidean distance calculations and linear mixed model analysis. For network analysis we used raw filtered count tables, aggregated to the genus level as recommended by (Peschel et al., 2021). Due to their high dissimilarity, NP and CGP data were analyzed separately throughout this study, except when compared to each other.

### 5. Statistical analysis

Comparing the microbiome communities of NP and CGP was conducted at the genus level, following 5000 iterations of rarefaction and averaging genera counts. We performed CLR-correction to the counts and calculated multivariate dispersion based on Euclidean distance metric. Principal Component Analysis (PCA) was also based on Euclidean distances.

A Permutational Multivariate Analysis of Variance (PERMANOVA) test based on Euclidean and UniFrac distances was conducted using the terms: ploidy, precipitation,and location/ecotype (separately for NP/CGP, respectively). Shannon diversity was analyzed for the rhizosphere bacterial communities in both NP and CGP samples grouped by precipitation ranges. Precipitation in NP and CGP was divided to three levels: high (>600 mm/year), mid (400-600 mm/year), and low (<400 mm/year). Taking the nested effect into consideration, we averaged diversity values per location/ecotype (NP/CGP, respectively) and used the calculated mean as a single replicate. For NP samples, the low precipitation level consisted of a single location and was therefore omitted. Wilcoxon test was done for the comparison between mid and high precipitation levels in NP; for CGP samples, statistical significance was first assessed using the Kruskal-Wallis test, followed by Dunn’s post-hoc test.

To identify ASVs unique to each precipitation level, ASVs were first filtered to retain those present in at least one sample per location and shared across all locations within a given precipitation level. Venn diagrams were then constructed to visualize precipitation-level–specific ASVs. The MaAsLin2 package was applied on clr-corrected rarefied counts to identify ASVs that were significantly associated with different locations/ecotypes using the three locations/ecotypes that fall within the mid-precipitation range as reference. A significance threshold was applied (adjusted P ≤ 0.05).

Normalized stochastic ratios (NST) were calculated using the package NST (v.3.1.10, (Ning et al., 2019)). We combined the two datasets (NP and CGP) to calculate NSTs per dataset and NST per location. To find NST per precipitation level we used the separate datasets after averaging the replicates per location. All NST values were based on Jaccard distances calculated from filtered raw ASV counts.

Network comparison analysis was conducted using the NetCoMi package (Peschel et al., 2021). This package allows a construction of two networks in parallel, comparing their different characteristics. We compared the networks of high and mid precipitation levels within each experiment, *i*.*e*., NP and CGP. The construction of the various networks was based on selecting the 200 most frequent genera in both datasets, using raw filtered genera count tables, applying the spiec-easi algorithm with internal clr-transformation. Hub taxa were identified using Eigen centrality, and clusters were computed using the Louvain algorithm. Network pair-wise comparisons were conducted by applying randomized permutations (n=100), significance was determined based on Benjamini-Hochberg corrected p-values. All data visualization and manipulation were conducted using the R packages vegan (Oksanen et al., 2007), phyloseq (McMurdie & Holmes, 2013), ggplot2 (Wickham, 2016), ggpubr (Kassambara, 2020), ggsci, vennDiagram (Chen & Boutros, 2011), and tidyverse (Wickham et al., 2019).

## Results

The bacterial community compositions of the rhizosphere microbiome of *Brachypodium* plants in Natural Populations (NP) and in a pot experiment (Common Garden Populations - CGP) were mostly represented by Actinomycetota and Pseudomonadota, respectively (Fig. S2). Principal component analysis revealed marked differences in within-population microbiome structure between NP and CGP (Fig. S3). Microbiomes associated with NP plants exhibited a higher variance between community compositions of the different samples, as observed by PCA and by multivariant dispersion values, 21.29 and 14.83 for NP and CGP, respectively (ANOVA p<0.001). However, despite this overall dispersion, samples collected from the same location consistently clustered together, indicating reduced microbiome distances within sites relative to between sites. These patterns confirm that environmental heterogeneity in natural settings promotes divergent microbiome assembly, whereas standardized conditions in the common garden constrain variability and homogenize microbiome structure. Accounting for the location of sample collection (NP) or ecotype (location of origin seed collection, CGP) significantly explained most of the microbial community variance (49.8% and 50.4% for NP and CGP, respectively, based on Weighted UniFrac distances), after accounting for plant ploidy (Table 1). Although NP and CGP differed in their degree of ordination dispersion, PERMANOVA indicated that sample origin explained a similar proportion of variation in both populations. This suggests that differences among origins are robust but are expressed against contrasting levels of within-population heterogeneity.

**Table 1.** Permutational multivariate analysis of variance (PERMANOVA) table using weighted Unifrac distances showing the variance explained by plant ploidy and location/ecotype on the rhizosphere bacterial communities of NP and CGP samples, respectively. In CGP, ecotype refers to the original location from which the seeds of the ancestral plants were collected.

| Data set | Variable | R<sup>2</sup> | P |
| --- | --- | --- | --- |
| NP | Plant ploidy | 0.021 | 0.560 |
|  | Location | 0.498 | 0.001 |
| CGP | Plant ploidy | 0.095 | 0.003 |
|  | Ecotype | 0.504 | 0.001 |

To evaluate bacterial alpha diversity, we calculated Shannon and observed diversity indices for both NP and CGP samples (Fig. S4). In both cases, no statistically significant differences in diversity were detected across locations (p = 0.19) or ecotypes (p = 0.075). A higher Shannon diversity was noticeable in CGP mid-precipitation locations (Ramat Avisar, Modiin, and Eshtaol Forest). Grouping the locations according to mean annual precipitation levels revealed that in CGP the different precipitation levels differed significantly in their microbial diversity (Figure 2), with mid precipitation level displaying higher diversity than high precipitation level. A similar trend was observed in NP, but was not found significant, possibly due to the low number of replicates.

**Figure 2.**
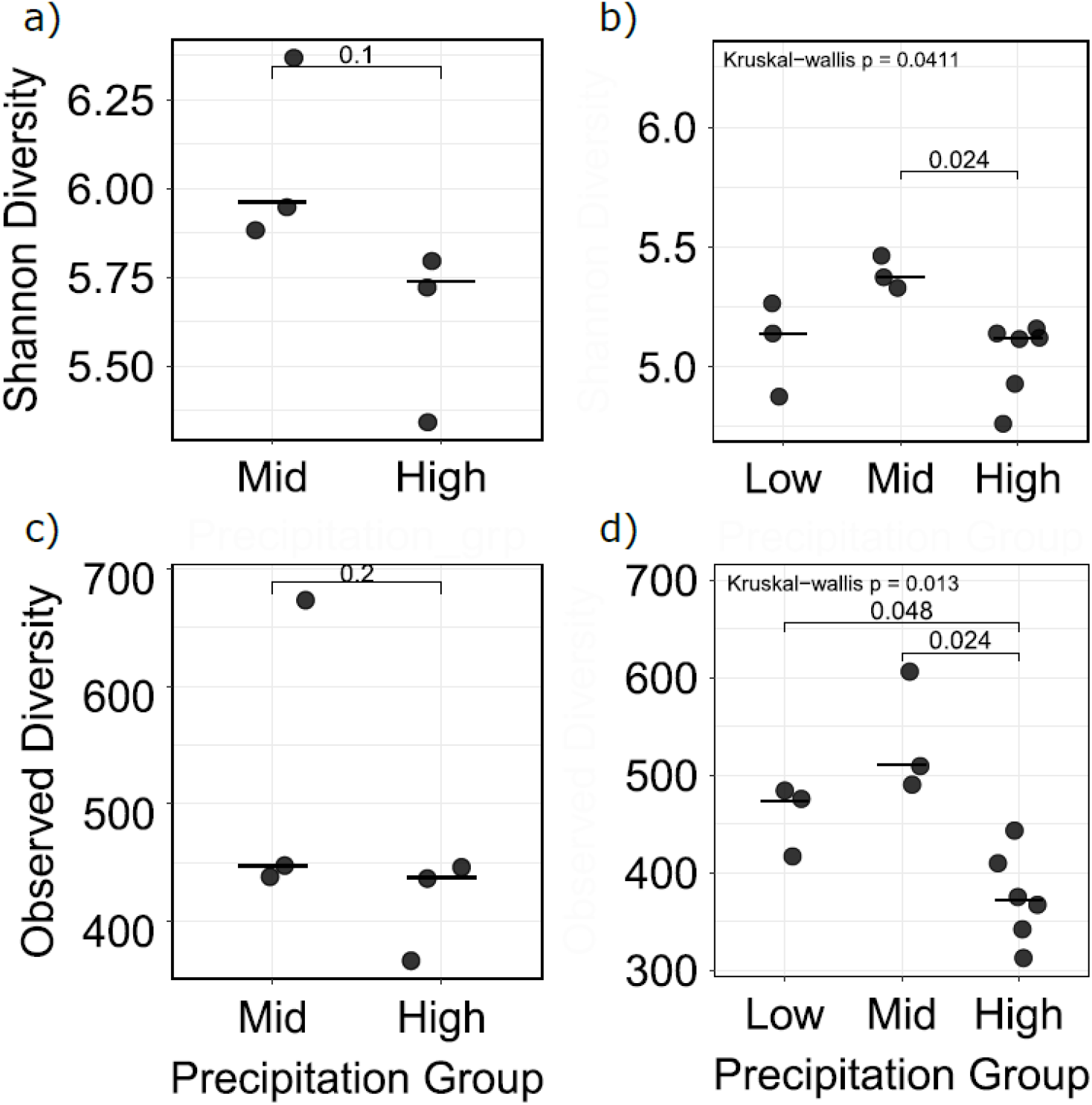
Diversity (a, b) and richness (c, d) of the microbiome of NP (a, c) and CGP (b, d) *Brachypodium* plants by precipitation level. Each dot represents the mean of all samples from a single location/ecotype.

We identified 152 and 81 unique ASVs in the mid and high precipitation levels of NP samples and 90, 110, and 7 ASVs unique to the low, mid, and high precipitation levels for CGP samples respectively (Figure 3). Overall, both NP and CGP samples showed a higher number of unique ASVs in the mid-precipitation level compared to the other precipitation levels. At the phylum level, unique NP ASVs were represented 12 phyla in the mid-precipitation level, compared with 10 phyla in the high-precipitation level; thus, the phyla Cyanobacteriota and Armatimonadota were uniquely represented in mid-precipitation level in NP (Figure 3). In CGP, unique ASVs represented 13, 12 and 6 phyla, for low, mid and high precipitation levels, respectively. Taxa uniquely present in the mid-precipitation level relative to the high-precipitation level included members of Bacillota (Planococaceae); Gemmatimonadota (Gemmatimonadaceae and Longimicrobiaceae); Planctomycetota(Rubinisphaeraceae, Gemmataceae, wd2101 soil group, and Pirellulaceae); Verrucomicrobiota (Pedosphaeraceae, Rubritaleaceae, Opitutaceae, and the class Verrucomicrobiae); and Myxococcota (Nannocystaceae, Haliangiaceae, Polyangiaceae, Blrii41, Sandaracinaceae, and the orders Bifdi19 and mle1-27).

**Figure 3.**
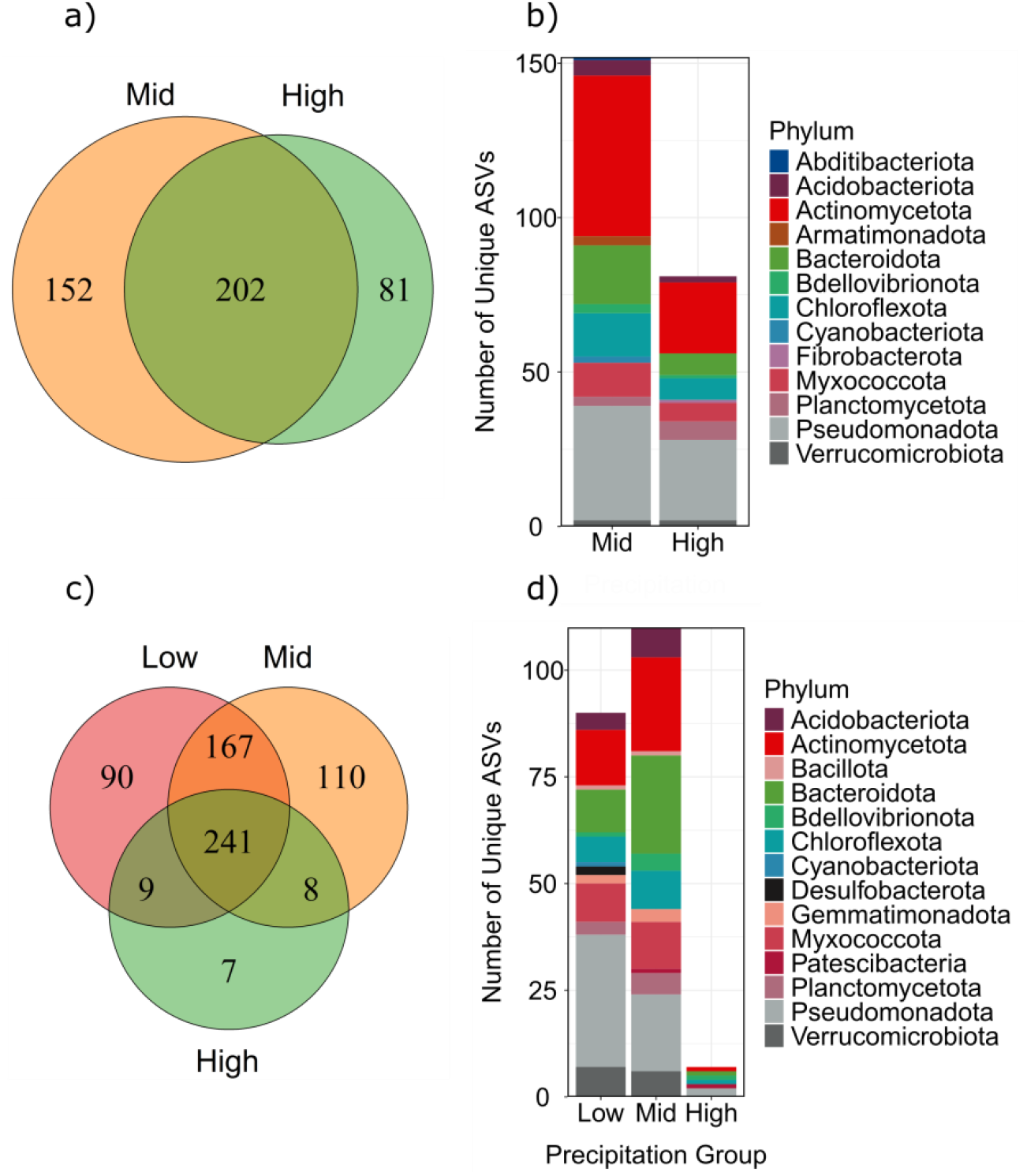
Venn diagrams displaying the number of ASVs shared among and unique to precipitation levels, considering only precipitation-level-specific ASVs within NP (a) and CGP (c). Bar plots represent the phylum-level taxonomic composition of precipitation-level–specific ASVs for NP (b) and CGP (d).

**Figure 4.**
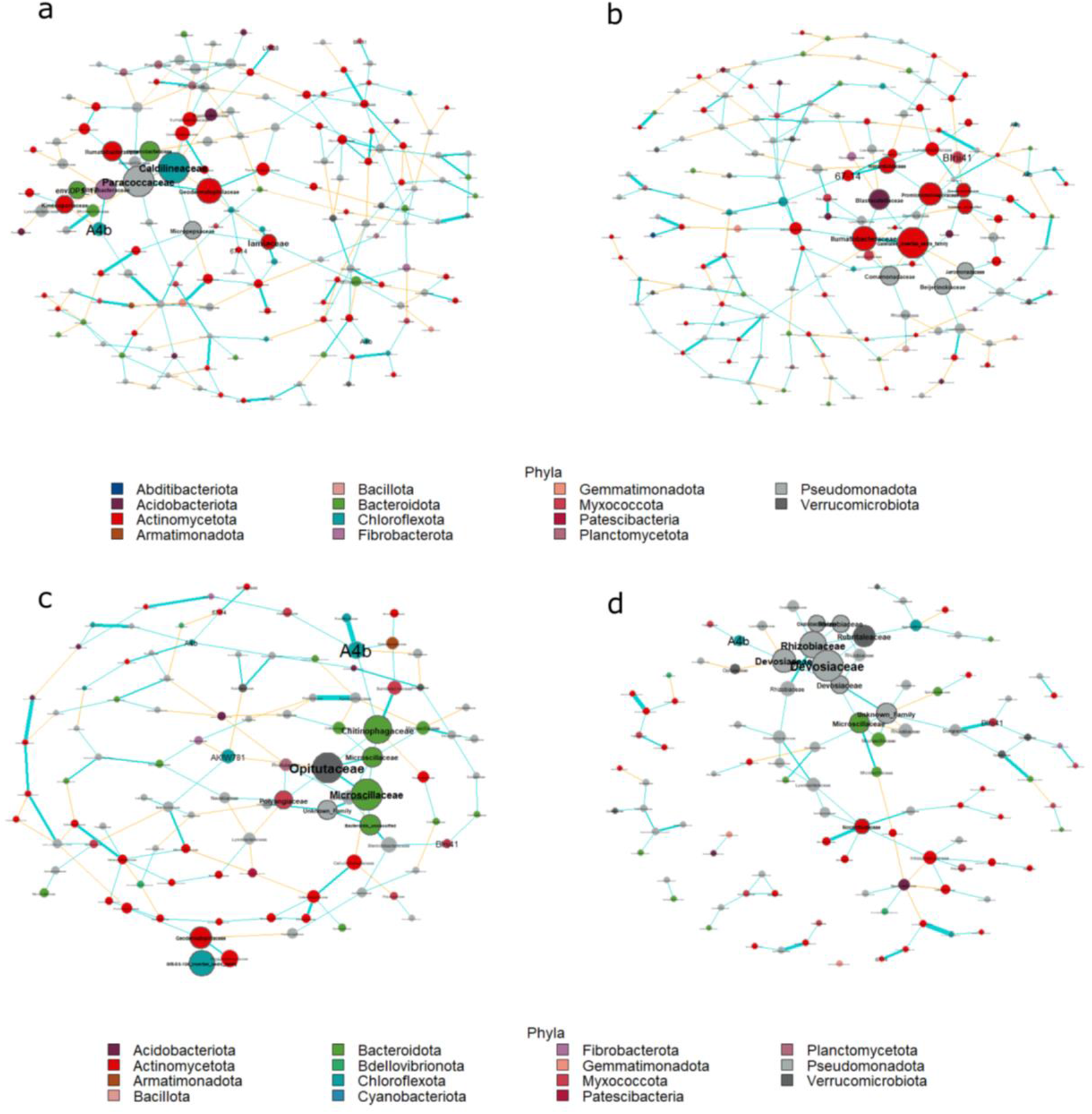
Comparison between mid (a, c) and high (b, d) precipitation levels of NP (a, b) and CGP (c, d). Each network displays the union of the 50 genera (nodes) with the highest closeness in both mid and high precipitation levels in either NP or CGP. Node size corresponds to the Eigen value of each node in each network, node colors correspond to the phylum, node outline represents hub taxa (dark – hub, light – non-hub), edge colors correspond to association type (blue – positive, yellow – negative), edge weights correspond to the relative strength of the association, thicker edges represent stronger association.

Since bacterial community composition was significantly associated with location/ecotype, and diversity showed an association with precipitation ranges in the sampling location or location of origin, we found ASVs that significantly associated with the different locations, using locations of the mid precipitation level as individual references. Overall, we found a total of 183 and 141 unique ASVs in NP and CGP, respectively, that significantly associated with either one of the high-precipitation locations/ecotypes compared with either one of the mid-precipitation locations/ecotypes; and 87 and 98 unique ASVs that associated with the mid-precipitation locations/ecotypes in NP and CGP, respectively (Table S2). We filtered the significantly associated ASVs to include only those that represent families that are consistently positively / negatively associated with the locations of the mid-precipitation level (Figure S5) and compared the results for NP and CGP. Among the families shared between NP and CGP locations/ecotypes associated with high precipitation, we found Streptomycetaceae, Nannocystaceae, Promicromonosporaceae, and Cellvibrionaceae. These families often inhabit soils and plants, degrade decaying plant matter, and secrete secondary metabolites (Cui et al., 2004; Kämpfer et al., 2014; Kohl et al., 1983; Tóth et al., 2021). Families that were significantly more abundant in locations/ecotypes of mid-range precipitation in CGP and NP included Cellulomonadaceae, Chitinophagaceae, Sphingobacteriaceae, and Roseiflexaceae These families occupy various environments, *e*.*g*., soil, plant roots, and aquatic environments; some can degrade complex polysaccharides such as cellulose and chitin, and some are associated with primary production and carbon cycling (Besaury et al., 2021; Figueiredo et al., 2022; Hanada, 2014; J. et al., 2025; Rosenberg, 2014; Stackebrandt et al., 2006).

A network analysis comparing mid and high precipitation levels in NP and in CGP revealed that in both datasets, the networks of mid-precipitation level significantly differed from those of high precipitation level in cluster membership, highest centrality, betweenness and closeness (Table S3). In NP, most of the hubs of the mid precipitation level were of the Actinomycetota phylum, while in CGP they were mostly of the Bacteroidota phylum (Table S4). In the high precipitation level Actinomycetota dominated the network hubs in NP, while in CGP network hubs Pseudomonadota were most prevalent. Actinomycetota likely acts as structurally important taxa in natural soils where fluctuating moisture and heterogeneous resources favor stress-tolerant, metabolically versatile organisms (Ebrahimi-Zarandi et al., 2023). In GCP, ecotypes from different arid regions favored taxonomically divergent hubs, indicating relaxation of environmental selection (such as drought) and increased fast-growing ecotype specific bacteria.

Normalized stochasticity ratio for NP and CGP showed a clear deterministic process in CGP compared with a stochastic process in NP, as expected from a controlled experiment compared with true environmental conditions (NST = 34.6% and 78.5% for CGP and NP, respectively, P < 0.001 by Wilcoxon test). When comparing NST between precipitation ranges per dataset, *i*.*e*., NP and CGP, we observed a general trend of significantly higher NST for the mid-precipitation range compared with the high precipitation range in both NP and CGP (Table 2).

**Table 2.** Values of normalized stochastic ratios per examined group. High, mid, and low refer to the precipitation level examined.

| Group | NST | P (Wilcoxon) |
| --- | --- | --- |
| NP | 0.785 | <0.001 |
| CGP | 0.346 |  |
| NP-High | 0.756 | <0.001 |
| NP-Mid | 0.822 |  |
| CGP-High | 0.646 | <0.001 |
| CGP-Mid | 0.885 |  |
| CGP-Low | 0.774 |  |

## Discussion

Copious studies have established the difference between the microbiomes of plant rhizosphere and of the bulk soil, merely centimeters away from the root. Plant activity, exudates and physical properties create a unique set of conditions that appeal to a subset of the soil bacteria, creating a distinct microbiome that is unique to plant species, and is affected by plant parameters such as developmental stage, ploidy and genome structure (Anneberg et al., 2023; Gat et al., 2025; Ji et al., 2023). Evidence is increasing that plants have evolved to manipulate the rhizosphere, mostly via secretion of secondary metabolites, specifically to draw beneficial bacteria, while bacteria evolved to harness these exudates (Haney et al., 2015; Kazarina et al., 2025). This plant-microbiome co-adaptation constitutes a “holobiont” in which the plant microbiome is an extension of the plant phenotypes. This study compares the microbiome of *Brachypodium* rhizosphere in wild plants collected from various locations and in common-garden plants descending from wild plants collected from various locations across the aridity gradient in Israel.

*Brachypodium* species exhibit strong biogeographic structuring across the circum-Mediterranean region, with closely related taxa occupying distinct climatic niches along gradients of temperature and aridity (López-Alvarez et al., 2015). Across this range, populations show clear signatures of aridity-driven adaptation, including shifts in phenology, life-history strategies, and genomic regions associated with drought tolerance, particularly in the more arid-adapted allopolyploid *B. hybridum* (Wang et al., 2025). Host genotype in *Brachypodium* plays a significant role in structuring root and rhizosphere microbial communities, with distinct bacterial assemblages associated with different genetic plant lineages. Studies using diverse *B. distachyon* accessions have shown that genotype effects persist across soils and environments, indicating heritable control over microbiome assembly (Ricks et al., 2026). In our study, we observed a higher bacterial diversity in the rhizosphere microbiome of *Brachypodium* plants originally growing in sites with mid precipitation level, located in a semi-arid transition zone at the boundary of the Mediterranean climate, where annual precipitation decreases from 600 mm to 400 mm. The NP locations and CGP ecotypes were sampled from different climatic regions classified as arid (MAP < 400 mm/year); semi-arid (400-600 mm/year) and dry sub-humid (> 600 mm/year) (FAO, 2004). Thus, the locations represented in this study display a wide range of mean annual precipitation values, different climatic regions, and various edaphic conditions. These conditions were expected to affect the rhizobiome diversity and composition of a natural plant population, as was indeed observed. Moreover, the alpha diversity of the rhizosphere bacterial communities of plants grown in a common garden also exhibited correlation with these conditions. A previous study using a different set of *Brachypodium* plants reported a lower but significant genotype effect, explaining ∼23% of bacterial community variation (Ricks et al., 2026). Notably, that study focused exclusively on *B. distachyon*, whereas our CGP experiment included multiple genetically and ecologically distinct *Brachypodium* populations. Increased host genetic divergence is expected to amplify microbiome differentiation, potentially explaining the higher proportion of variance observed here.

As opposed to an increase in soil microbiome functional diversity that was demonstrated for a precipitation range of >600 mm (Tripathi et al., 2017), we observed that for the mid-precipitation level (400-600 mm) bacterial diversity and richness in the rhizobiome were significantly higher than in the high precipitation range. In addition, higher precipitation was also associated with a lower normalized stochasticity ratio and a higher number of positively associated ASVs. These observations suggest that plants representing ecotypes with higher annual precipitation require a more specific microbial community composition, thus applying a specific selection process that results in lower diversity and lower stochasticity. In contrast, higher microbial diversity, higher stochasticity, and lower number of associated ASVs in the mid precipitation level may suggest higher plasticity in these ecotypes. The decrease in precipitation level is a condition under which adaptation strategies may extend below ground, where hosts associate with diverse or functionally redundant microbial communities to buffer against fluctuating moisture and nutrient availability. The selection of a more diverse rhizosphere microbiome *Brachypodium* may allow for extended phenotypic plasticity.

Moreover, our results confirm that different plant ecotypes originating from contrasting climatic conditions, particularly along aridity gradients, differ in the microbes they recruit, confirming that plant adaptation to environment extends to below-ground biotic interactions. Bacterial community composition differed significantly according to the plant’s ecotype/location in the common garden and natural population, respectively. Similarly, (Kazarina et al., 2023) described an ecotype effect on the microbial diversity of plants in reciprocal gardens. When comparing the microbial community composition between the common garden and the natural population, a clear difference was observed in beta-diversity, differentially abundant bacteria and network compositions, per location or precipitation range. We suggest that the different taxonomic composition stems from the differences in soil bacterial composition between these populations. Among the ASVs that were significantly associated with the different locations/ecotypes, we observed similar patterns according to precipitation ranges. High-precipitation range was characterized by fewer associated ASVs of the Alphaproteobacteria, Bacteroidia, and Blastocellia classes, while ASVs of the Actinobacteria class were more commonly associated with high precipitation range in NP and CGP. In addition, network hubs of the high-precipitation level in both NP and CGP displayed lower phyla diversity, as compared with mid-precipitation level. These similarities in association patterns suggest similarity in microbial recruitment or co-adaptation. Examining the functional profiles of these communities should shed more light on the suggested co-adaptation. Deciphering the interactions between *Brachypodium* plants and their microbiome over an aridity gradient could provide insights on the other members of the Poaceae family, specifically wheat, and their adaptation to a changing climate.

