## Supplemental Information for "Rainfall legacy effects on the rhizosphere bacterial diversity of *Brachypodium* ecotypes across an aridity gradient"

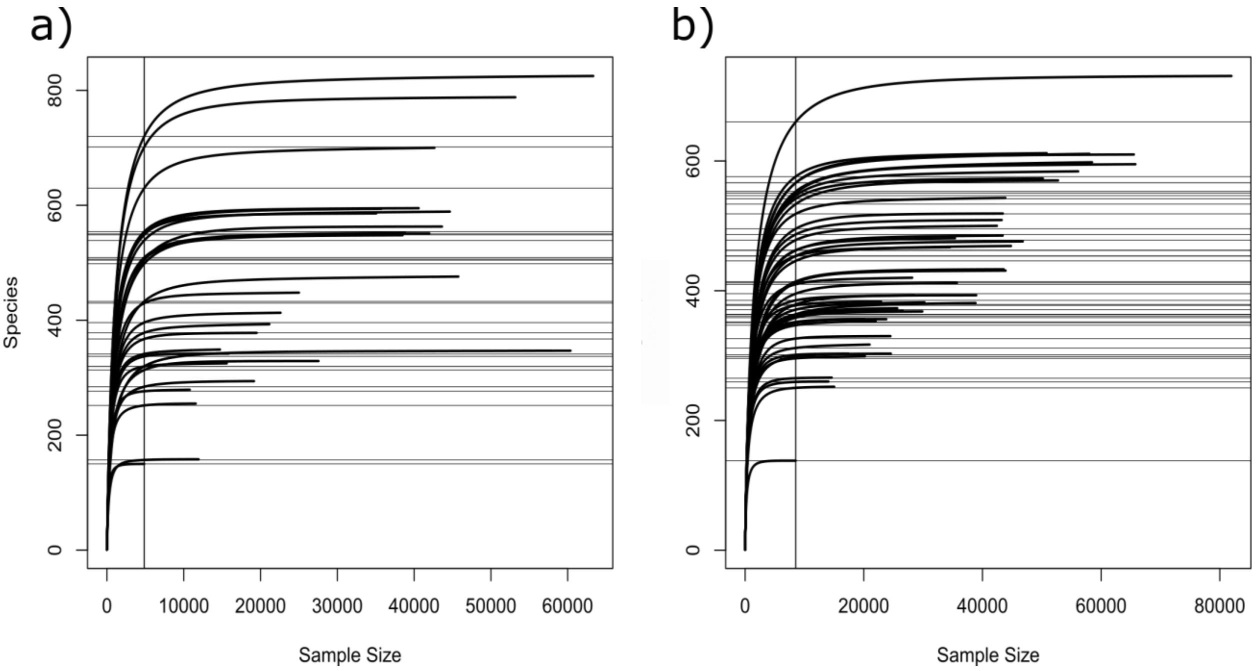


Figure S1: Rarefaction curves for a) NP and b) CGP samples for amplicon sequencing data. Rarefaction curves depict the relationship between sequencing depth (x-axis) and observed ASV richness (y-axis) per sample.

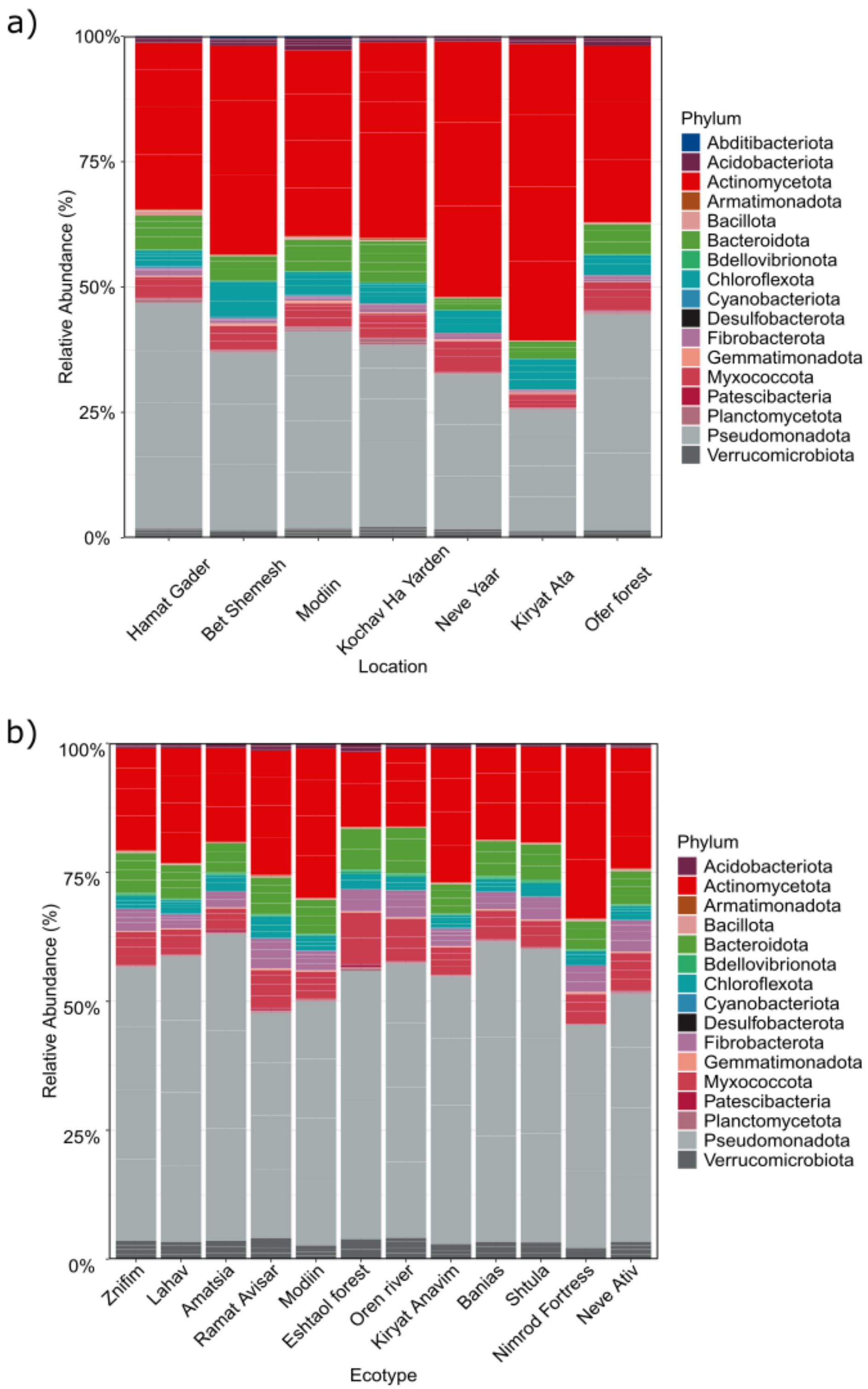


Figure S2: Taxonomic composition of NP (a) and CGP (b) by relative abundance of phyla.

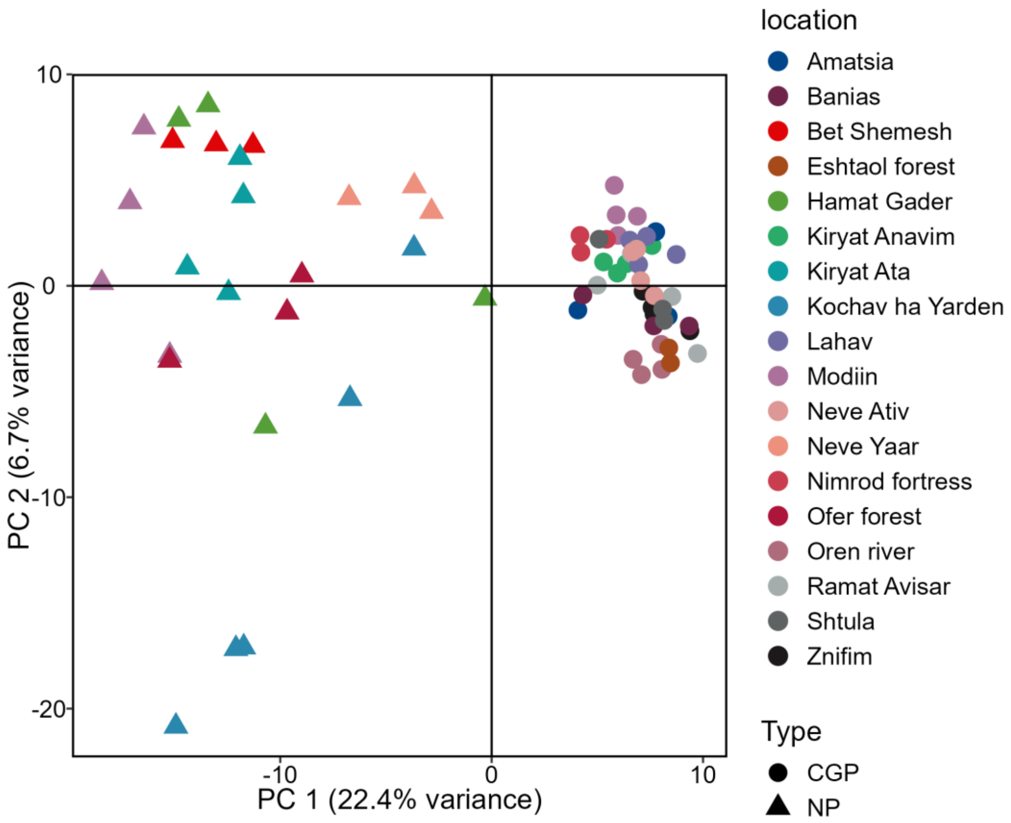


Figure S3: PCA based on Euclidean distances of NP and CGP *Brachypodium* rhizobiome.

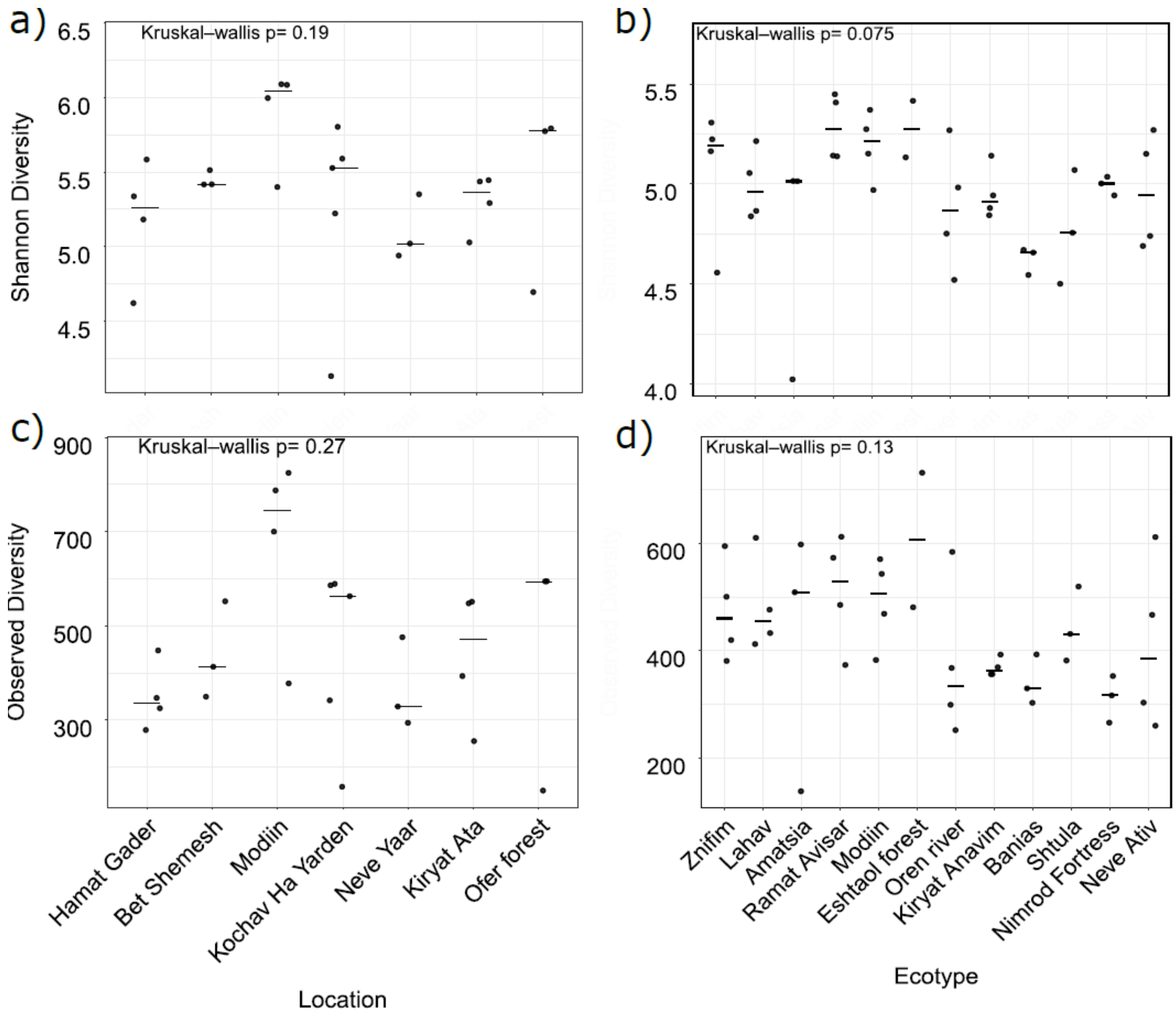


Figure S4: Shannon (a, b) and observed (c, d) diversity and richness indices of NP (a, c) and CGP (b, d) of *Brachypodium* rhizobiome by plant location / ecotype for NP and CGP, respectively.

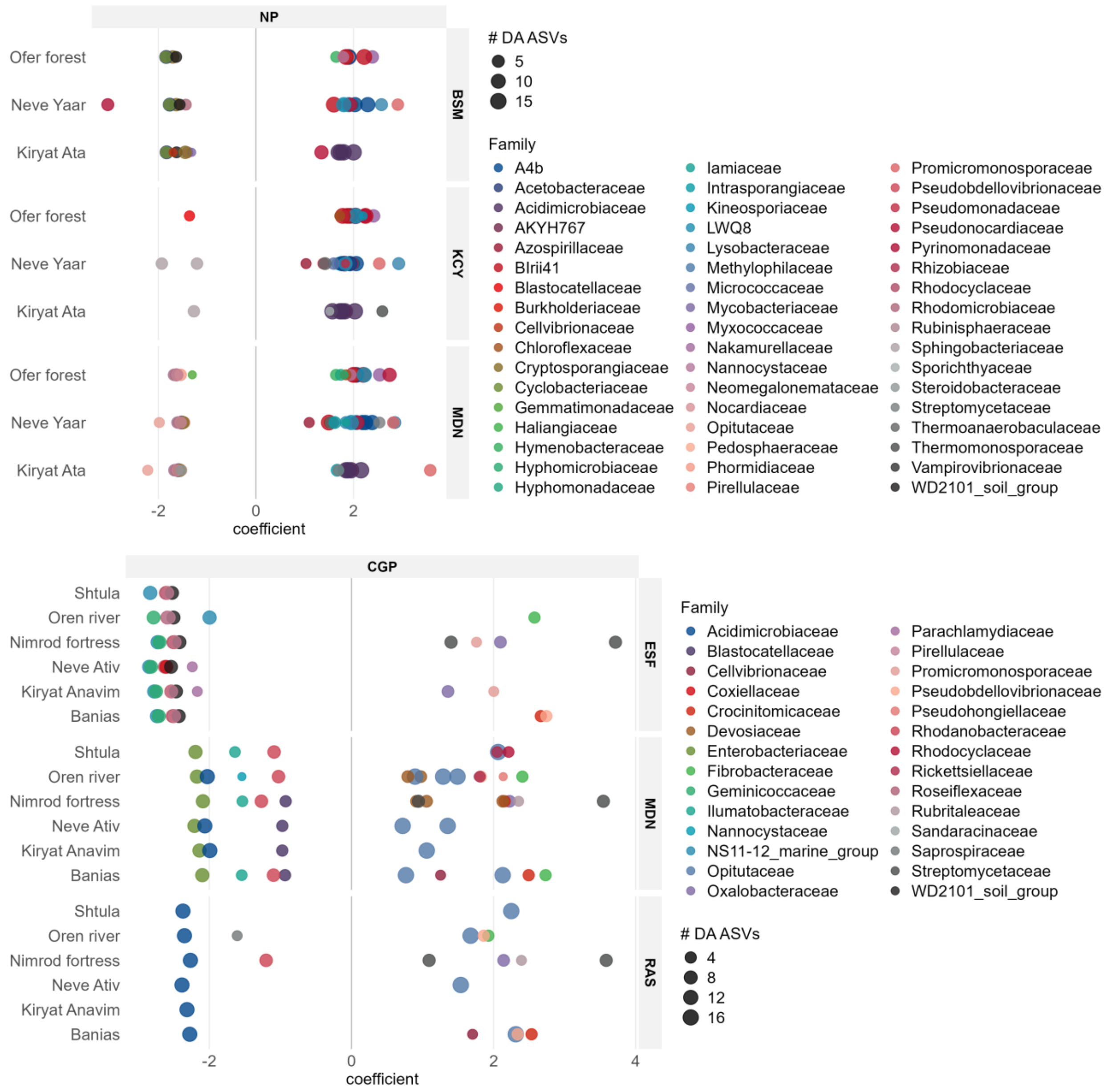


Figure S5: Taxonomic families comprised of ASVs that were significantly associated with either mid-precipitation locations (negative coefficient) or high precipitation locations (positive coefficient) in NP (upper) and CGP (lower). Symbols' color represents taxonomic family; symbol size represents the maximal number of individual ASVs that were significantly associated with any location per family. Labels on the right are mid-precipitation level location/ecotype abbreviations: BSM – Beit-Shemesh; KCY – Kochav Ha Yarden; MDN – Modiin; ESF – Eshtaol forest; RAS – Ramat Avisar.
